# The self-regulated response of the Wnt pathway to an oncogenic mutation in β-catenin

**DOI:** 10.1101/2021.05.05.442673

**Authors:** Ana R Moshkovsky, Wenzhe Ma, Marc W Kirschner

## Abstract

Oncogenic mutations in β-catenin can inhibit degradation of β-catenin by preventing phosphorylation of its degron. Degron phosphorylation is mediated by the Axin scaffold, Casein Kinase 1α (CK1α) and Glycogen Synthase Kinase 3 (GSK3). We studied an oncogenic form of β-catenin with a deletion of serine 45 (S45), a site that is phosphorylated by the kinase CK1α. When the S45 site is phosphorylated, it promotes the GSK3-mediated phosphorylations of the β-catenin degron. Deletion of S45 would be expected to prevent GSK3-mediated phosphorylation of the mutant protein and thus block degradation. We found that the S45 mutant was still phosphorylatable by GSK3, and its expression increased the concentration of Axin, restoring the rate of GSK3-mediated phosphorylation to levels comparable to those observed for the wild-type β-catenin. We conclude that there is one core mechanism for creating the phosphodegron for both primed and unprimed β-catenin, which involves the generation of an Axin-GSK3 complex.

**SIGNIFICANCE:** Understanding how the Wnt pathway responds to mutations in β-catenin phosphodegron can reveal important properties of the pathway in normal and cancer cells and is valuable for the design of more effective therapeutic strategies.

## INTRODUCTION

An understanding of the function and action of cellular components has been greatly facilitated by naturally occurring genetic mutations and by genetic mutations generated experimentally. However, in many biological systems, and perhaps in most biological systems, the gene products participate in complex pathways involving multiple levels of controls and complex feedback circuits. In such situations, the extrapolation from mutation to mechanism often is fraught with ad hoc assumptions that can lead to erroneous conclusions, even when the basic observations are sound. The potential for such misunderstanding is especially severe in the Wnt pathway which depends on transcriptional control and feedback, control of protein stability, posttranslational modification, and subcellular localization not only of the principal target, β-catenin, but some of the regulatory components, principally but not limited to the scaffold proteins Axin1 and Axin2 and also of the Frizzled Wnt receptors (1). The importance of understanding the mechanism could be critical in the design of therapeutic agents that target the Wnt pathway.

The Wnt pathway was discovered in flies, frogs and mammalian systems about 40 years ago and was quickly recognized to be important in embryonic development and tumorigenesis. The secreted Wnt ligands of what is now known as the canonical Wnt pathway bind to transmembrane receptors, and signal through a still poorly understood pathway causing a rise in β-catenin, activating the transcription of specific genes, whose promoter has binding sites for TCF-β-catenin (1). Numerous studies have shown that binding of Wnt proteins to the receptor acts as an inhibitory signal, blocking constitutive degradation of β-catenin. As the Wnt ligands generally do not affect the rate of β-catenin synthesis (2), it is the inhibition of degradation that leads to a rise in β-catenin level; this in turn activates transcription at specific sites on the genome. The formation of a phosphodegron is the key and obligate step in the pathway of degradation, which is indirectly antagonized by the Wnt signal. The phosphodegron is generated by two kinases, casein kinase 1 alpha (CK1α), and glycogen synthase kinase (GSK3). CK1α phosphorylates β-catenin at serine 45 (S45) on β-catenin and this phosphorylation is required for the subsequent phosphorylations by GSK3. GSK3 then phosphorylates threonine 41 (T41), serine 37 (S37) and serine 33 (S33) (3, 4). As a result of these phosphorylations, a binding site is created for an E3 ligase, β-TRCP, which transfers ubiquitin to β-catenin, is recognized and degraded by the proteasome. Wnt activation partially inhibits β-catenin degron phosphorylation (2), leading to the accumulation of β-catenin. The current model is that there is an obligate process of priming phosphorylation by CK1α for the phosphorylations of the three downstream GSK3 sites to occur (3, 4).

The effect of mutations in β-catenin degron was studied by Vogelstein and Kinzler in HCT116 cells (5). HCT116 is a human colorectal cancer cell line that has a wild-type (WT) allele and a mutant allele of β-catenin with a full deletion of the codon encoding the serine 45 priming site in one allele (WT/ΔS45). Therefore, HCT116 cells co-express the WT and a mutant form of β-catenin. To study the importance of priming phosphorylation in β-catenin regulation, two cell lines were engineered from the HCT116 cells by disrupting either the mutant (WT/-) or the WT (-/ΔS45) allele of β-catenin (5). The expectation was that the mutant lacking the S45 phosphorylation would not bind GSK3 and therefore, be defective in the GSK3 mediated downstream phosphorylation of β-catenin. Since the mutant would fail to be degraded it would accumulate. Surprisingly, Vogelstein and Kinzler found that the mutant form lacking the priming site was still phosphorylated on the sites that are normally phosphorylated by GSK3 and that the level of phosphorylation was nearly as great as in the unstimulated condition (6, 10). This unexpected finding seemed only explainable by the involvement of other unknown components or undiscovered properties of the system. Vogelstein and Kinzler concluded that in the tumor cancer cells, mutant β-catenin must be modified by a phosphorylation process different from that used in normal tissues, which could open up other approaches to pharmacology of the Wnt pathway in tumor cells. It is worth considering, however, whether there is a different explanation that does not invoke new pathologies but rather depends on normal features of the Wnt pathway

In considering these results the key question is to what degree does obligate priming at S45 play in the degradation of β-catenin. If it were required, how would we explain the measurements that show that S45 mutations still lead to high levels of downstream phosphorylation. If under some circumstances it played no role, what are those circumstances? The resolution of this enigma comes from the fact that β-catenin does not act alone. The dynamically balanced Wnt pathway, comprised of several essential gene products, can be pushed into a different state using the same core mechanism that normally regulates WT β-catenin degradation. Hence there is no need to postulate a shadow pathway that comes into play when S45 is deleted.

Here we confirm that GSK3 is indeed able to phosphorylate mutant β-catenin (-/ΔS45), demonstrating that there is a potential direct path to GSK3-dependent phosphorylation for unprimed β-catenin. We show that to understand how in the wild type this does not occur to a high level, yet in the knockout mutant it does, we need to consider in detail the dynamic equilibrium between β-catenin and its negative regulators in terms of a simple model of mass action. This shift in equilibrium can be understood in terms of the amplified effects of a small increase in the level of β-catenin on the levels of Axin1 and Axin2. Together, this perturbation of Axin1 and Axin2 levels restores the rate of GSK3-dependent phosphorylation by mass action to levels comparable to that seen for WT/- β-catenin.

## RESULTS

### ΔS45 β-catenin can be phosphorylated by GSK3

It was previously shown that mutant ΔS45 β-catenin, mutated at the CK1α priming site for downstream GSK3 phosphorylation, is nevertheless phosphorylated (6) at the sites T41/S37/S33. Here, we not only confirm that ΔS45 β-catenin is phosphorylated at T41/S37/S33, as detected by a phospho T41/S37/S33 β-catenin antibody, but we also indicate that GSK3 is the agent for phosphorylation, as lithium chloride (LiCl), a specific inhibitor of GSK3, strongly inhibits the T41/S37/S33 phosphorylation of ΔS45 β-catenin (fig. 1).

**Figure 1.**
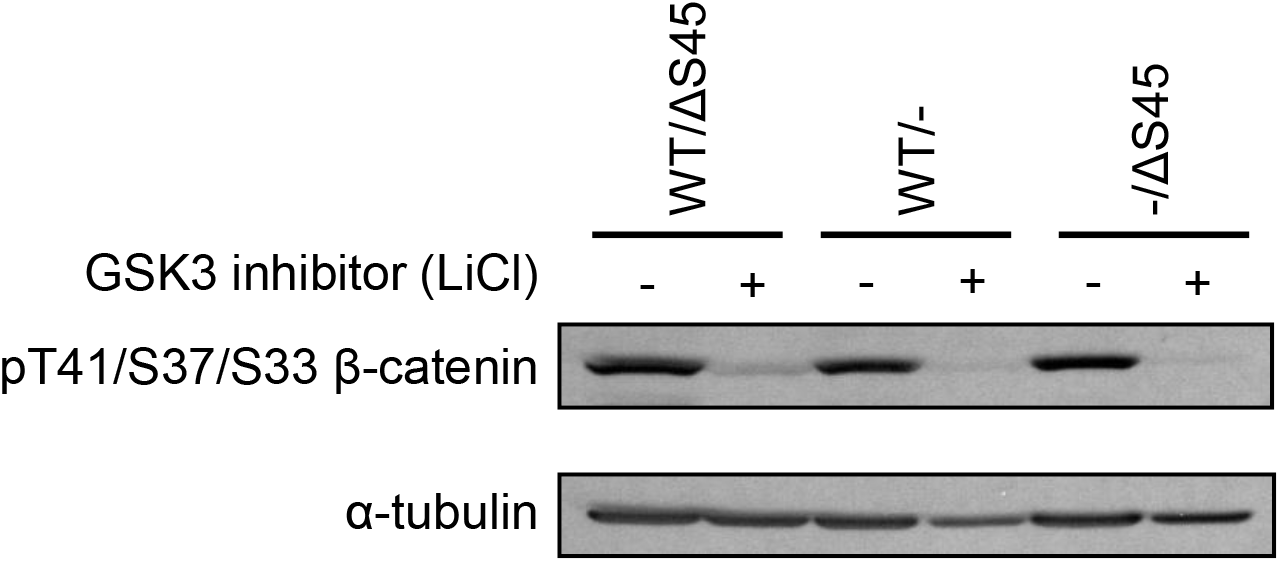
β-catenin is phosphorylated at T41/S37/S33 by GSK3 in WT/ΔS45, WT/- and -/ΔS45 β-catenin cells. Treatment with 40mM lithium chloride (LiCl) for one hour inhibited GSK3 phosphorylation at T41/S37/S33 in these cells. α-tubulin is shown as a loading control.

### Axin1 and Axin2 proteins increase markedly in -/ΔS45 β-catenin cells

The expression of Axin is key to explaining the S45–independent phosphorylation of ΔS45 β-catenin by GSK3. In vitro, Axin causes an increase in the rate of GSK3-dependent phosphorylation for unprimed β-catenin of as much as 20,000 fold, while S45 priming phosphorylation by CK1α increases phosphorylation by 30 fold (7). We therefore measured Axin1 and Axin2 protein levels in mutant ΔS45 and wild-type WT cells. We found that -/ΔS45 cells express 2.7− and 11.4-fold higher concentrations of Axin1 and Axin2 proteins, respectively, compared to WT/- cells (fig. 2A and 2B). The high Axin2 protein levels in -/ΔS45 cells is caused by higher Axin2 mRNA expression, which is 9.7 fold greater than in WT/- cells (Fig 2C). Presumably, the high Axin2 transcription is a result of higher levels of ΔS45 β-catenin, as Axin2 is a well-known Wnt/β-catenin transcriptional target. Axin2 mRNA and protein expression does not appear to have reached the maximum possible level in unstimulated -/ΔS45 cells since Axin2 mRNA and protein concentrations still respond to Wnt-3A stimulation in these cells (fig. 2C and 2D). Axin1 transcription is not regulated by Wnt/β-catenin and Axin1 mRNA levels are not significantly higher in -/ΔS45 cells than in WT/- cells (fig. 2C).

**Figure 2.**
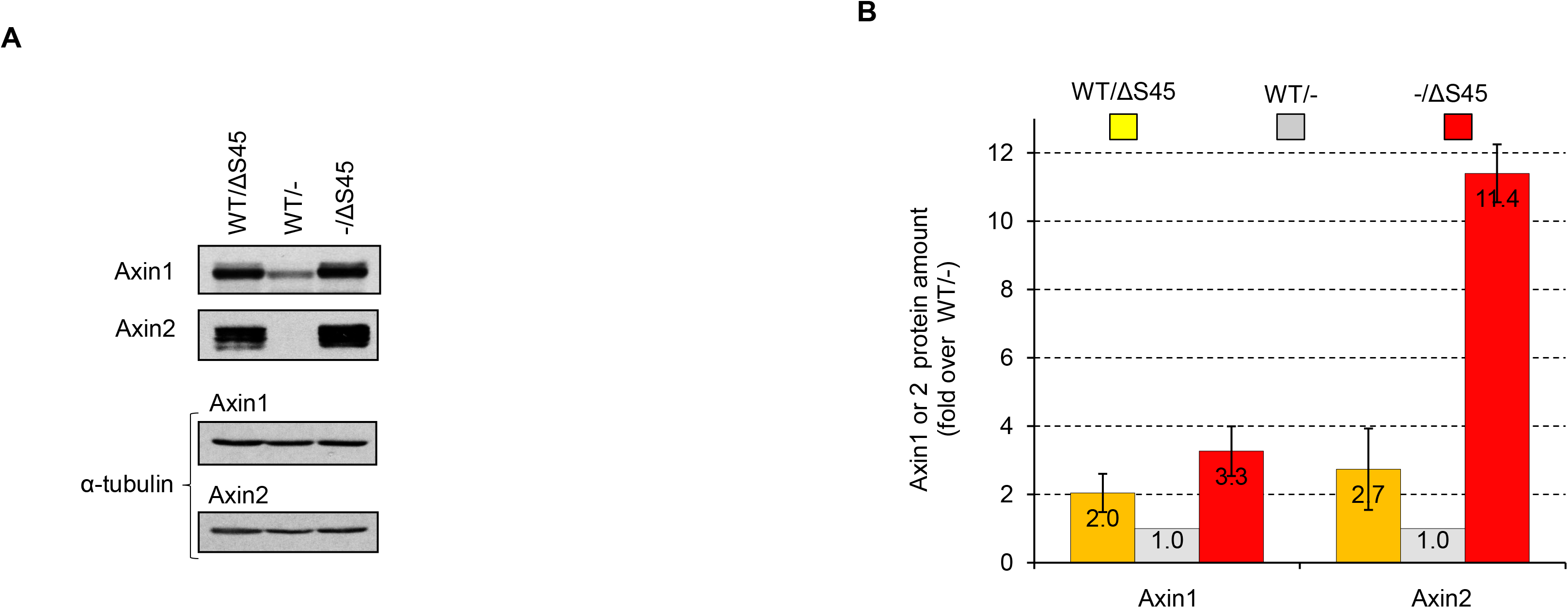
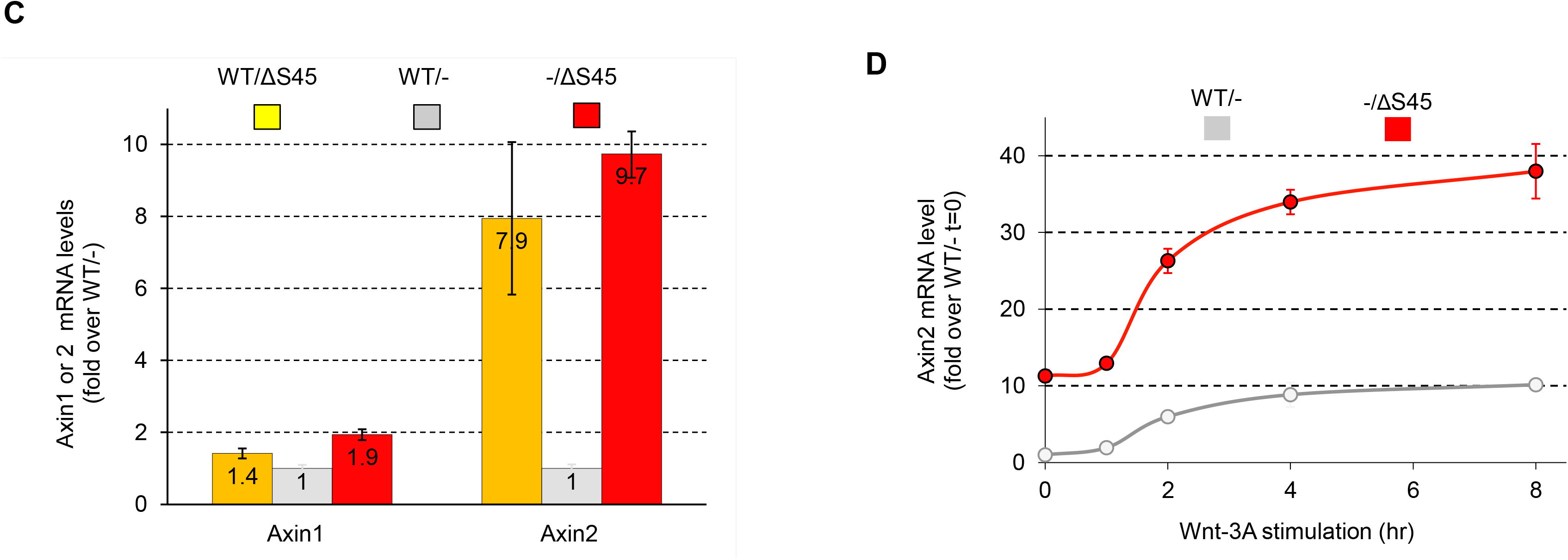
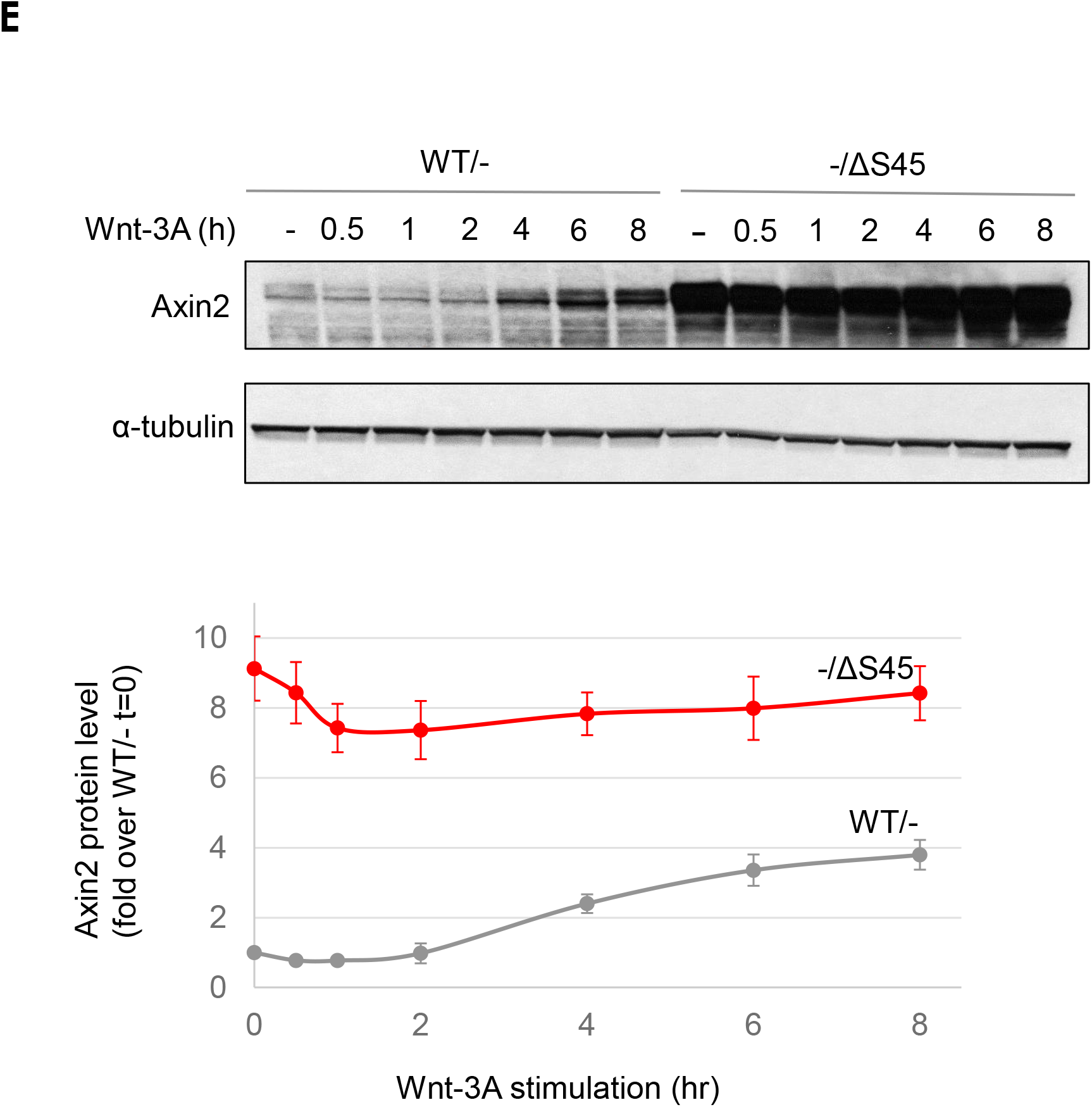
Axin1 and Axin2 concentration in cells expressing ΔS45 β-catenin. **A**, ΔS45 β-catenin expression increases Axin1 and Axin2 concentrations. **B**, Protein quantification of Axin1 and Axin2 in WT/ΔS45, WT/- and -/ΔS45 β-catenin <u>(n=3)</u>. **C**, ΔS45 β-catenin expression activates Axin2 transcription. **D**, -/ΔS45 cells show further activation of Axin2 transcription after Wnt-3A stimulation. **E**, Axin2 protein concentration increases in response to Wnt-3A stimulation in WT and ΔS45 cells.

In summary, -/ΔS45 cells respond to the mutational state of β-catenin through an increase in β-catenin, leading to increased levels of Axin1 and Axin2. Therefore, the priming phosphorylation of β-catenin by CK1α contributes to, but is not an absolute requirement for the subsequent GSK3-dependent phosphorylation and can be compensated by the rise of Axin1 and Axin2.

### Mass action and negative feedback enable ΔS45 β-catenin phosphorylation by GSK3

To elucidate the possible mechanisms that enable phosphorylation of ΔS45 β-catenin by GSK3, we analyzed the factors that affect the rate of the GSK3-mediated phosphorylation V^GSK3^ (V^GSK3^=k[β-catenin]). The two key factors are the concentration of β-catenin ([β-catenin]) and the rate constant *k* of GSK3-mediated phosphorylation.

One way to increase the rate of GSK3 phosphorylation V^GSK3^ by mass action is to increase the concentration of its substrate, β-catenin ([β-catenin]). Due to the more inefficient action of GSK3 without a priming phosphorylation, the level of mutant β-catenin accumulates to a higher level than the WT protein (fig. 3A). Higher amounts of β-catenin were also detected in the RKO human colorectal cancer cell line after silencing the expression of CK1α, the S45 priming kinase (fig. 3B). We conclude that though the priming phosphorylation at S45 significantly increases the rate of GSK3 phosphorylation, the rate in S45 mutants or deletions though lower, is not zero. As expected, the reduced rate of degradation of S45 mutants causes the level of β-catenin to rise, which by mass action increases the overall flux of degradation. In the absence of other effects, it will rise to a new stable level where the rate of degradation equals the rate of synthesis. Since there is no known effect on the rate of Wnt on β-catenin synthesis, this flux of β-catenin degradation should be the same as the wild type β-catenin. In other words, the S45 mutant on its own should cause the β-catenin to rise independently of Wnt addition.

**Figure 3.**
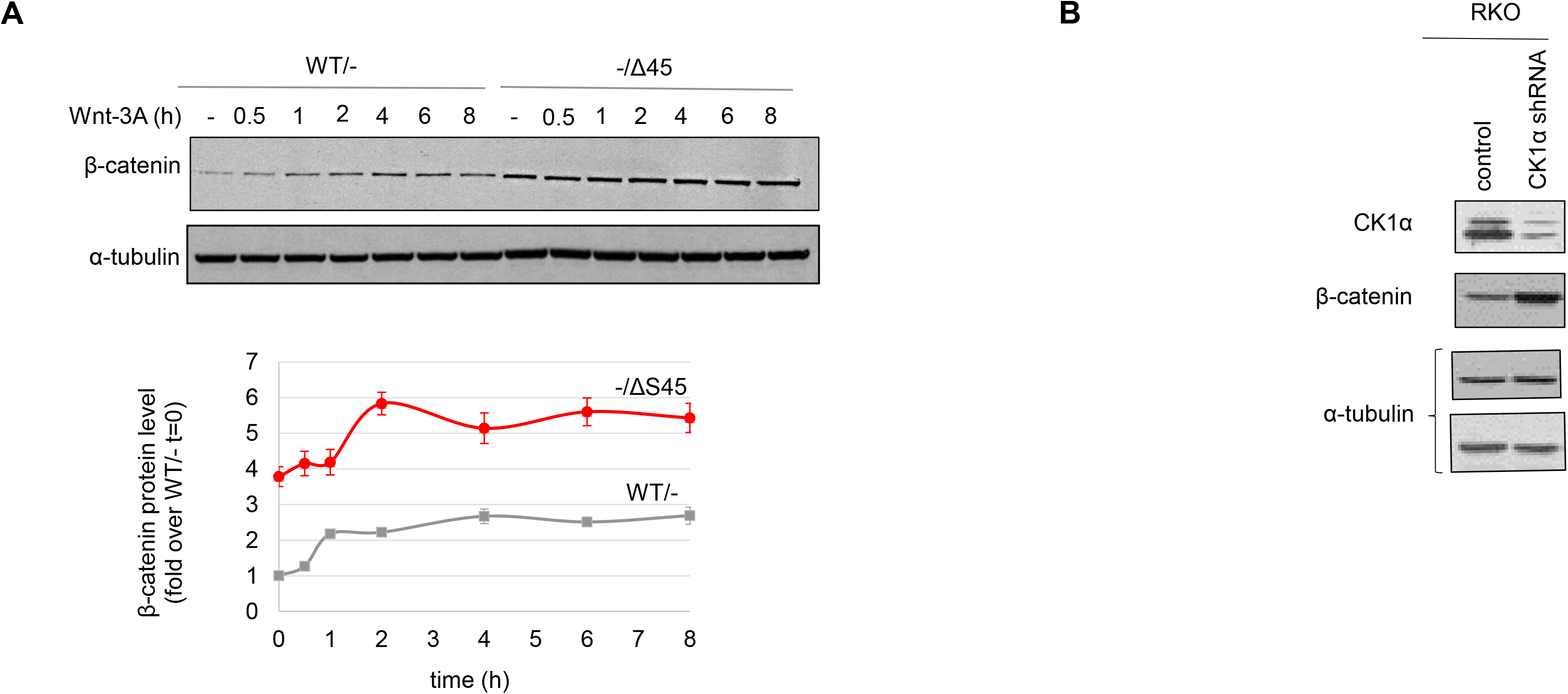
β-catenin protein expression in WT/- and -/ΔS45 cells before and after Wnt-3A stimulation. **A**, Western blotting of β-catenin expression and protein quantification of β-catenin in WT/- and -/ΔS45 cells (n=3). **B**, Western Blotting of β-catenin expression in RKO cells after CK1α silencing.

But how far will it rise? There is another effect tied to the increase in β-catenin level that opposes the rise in β-catenin and that is a rise in Axin levels. As the rate of GSK3-dependent phosphorylation is catalyzed by the Axin scaffolds, this increases the level of GSK3 mediated phosphorylation by increasing the catalytic rate for phosphorylation, simply by increasing the concentration of Axin-GSK3 complex.

As we demonstrated, the concentrations of both Axin1 and Axin2 increase in ΔS45 cells. The increased Axin concentrations, in turn, enhance GSK3-mediated phosphorylation, further compensating for the lack of priming S45 phosphorylation. As a result, T41/S37/S33 phosphorylation in mutant cells reaches a concentration comparable to that in WT cells (fig. 2).

To test the hypothesis that the feedback generated by increased Axin expression causes the increased ΔS45 β-catenin instability, we compared the amount of WT β-catenin in WT/- and WT/ΔS45 cells. If WT-β-catenin and ΔS45 β-catenin expression in the same cell does not change the stability WT β-catenin, then this would suggest that there is no negative feedback through increased Axin1 and Axin2 expression (fig. 4A). Alternatively, if ΔS45 and WT β-catenin co-expression decreases the stability of WT β-catenin, then it indicates the existence of a negative feedback, such as that caused by increased Axin1 and Axin2 (fig. 4A). To distinguish the WT β-catenin from the ΔS45 β-catenin, we immunoblotted with an anti-phospho S45 antibody, which effectively distinguishes the two forms. The phospho specific S45 antibody does not cross react with ΔS45 β-catenin since ΔS45 β-catenin does not have that specific phosphorylation and lacks the S45 residue (fig. 4B and 4C). Using this specific phospho-S45 β-catenin antibody, we found that the amount of WT β-catenin is much higher in WT/- cells than in WT/ΔS45 cells indicating the presence of negative feedbacks in WT/ΔS45 cells (fig. 4D).

**Figure 4.**
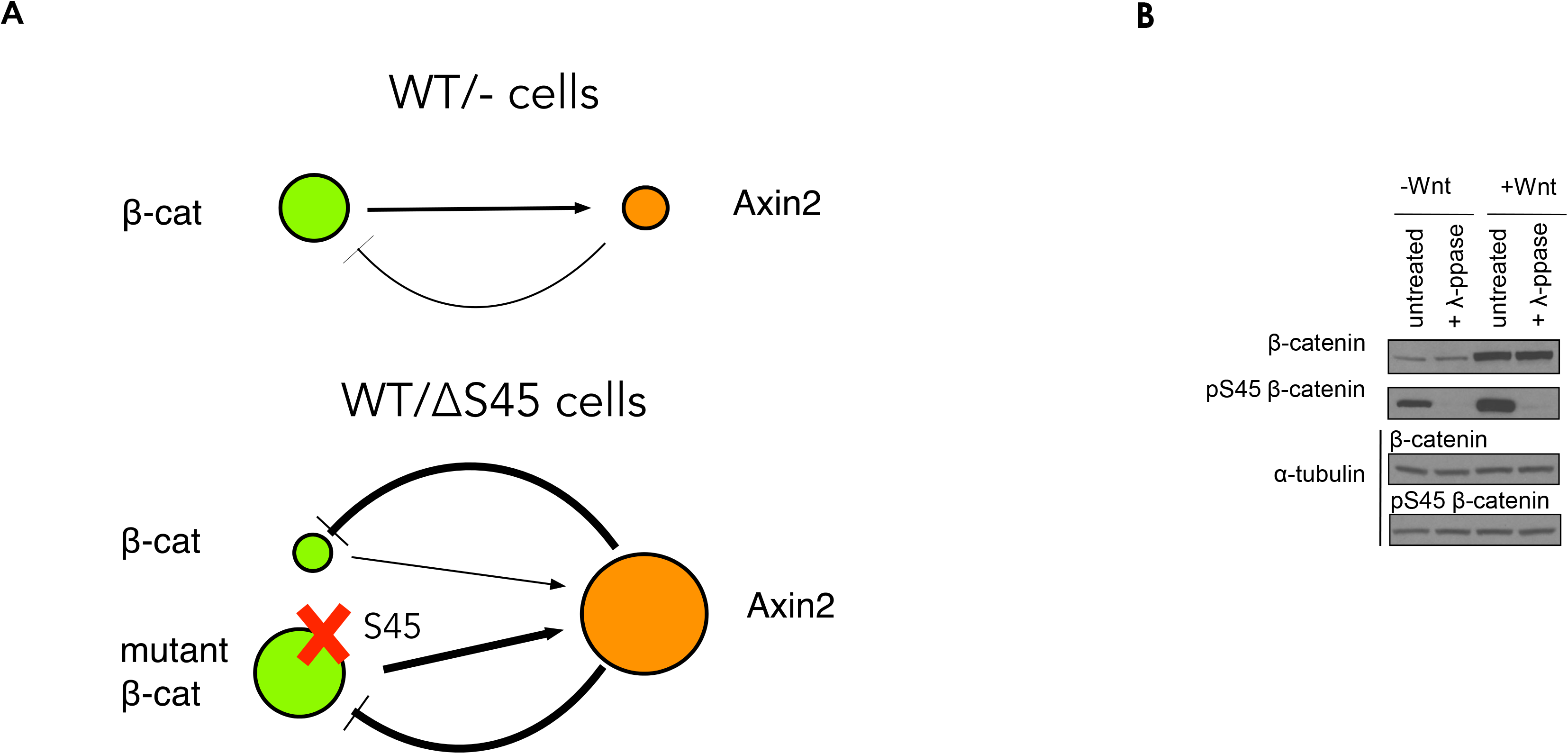
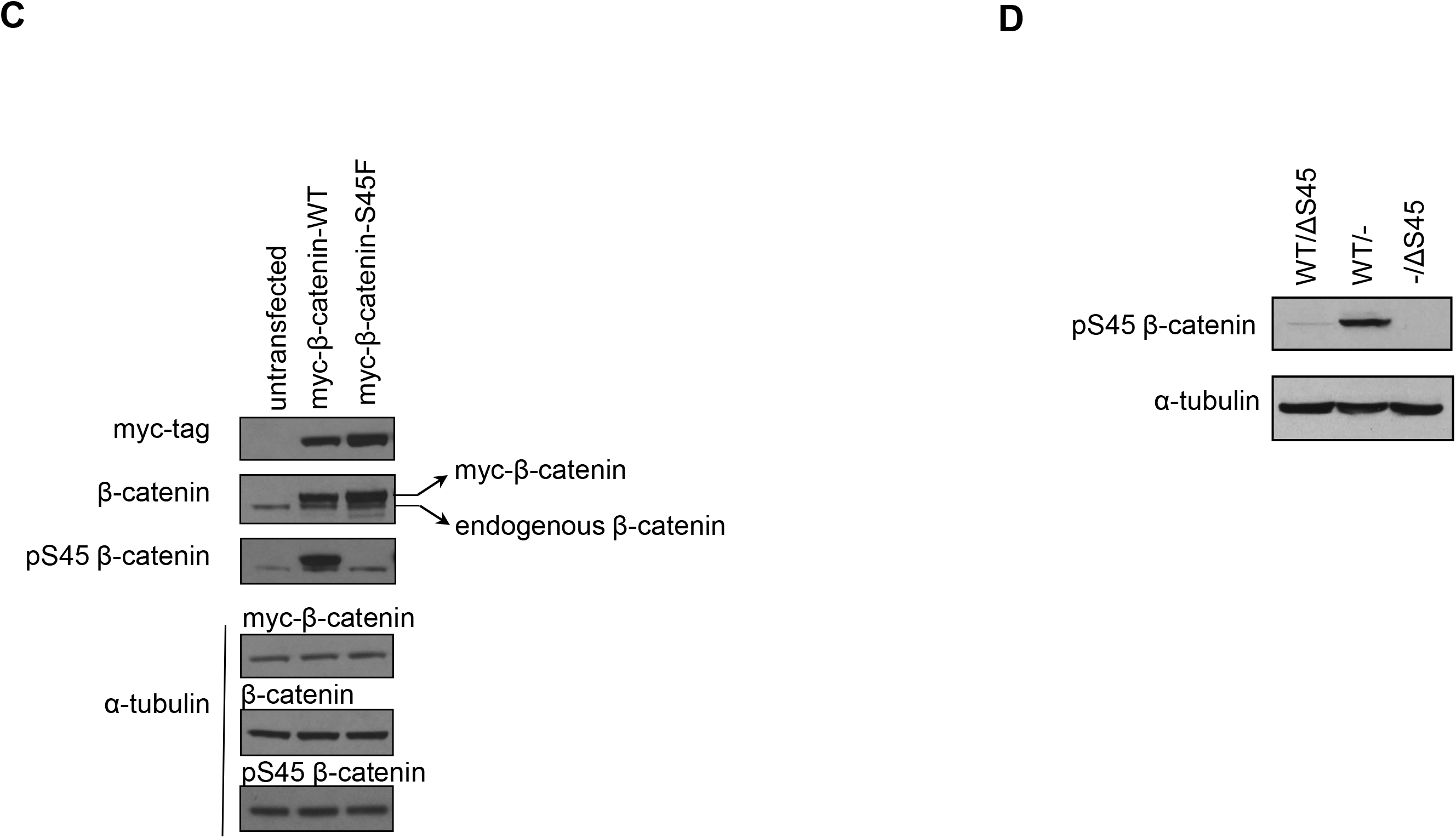
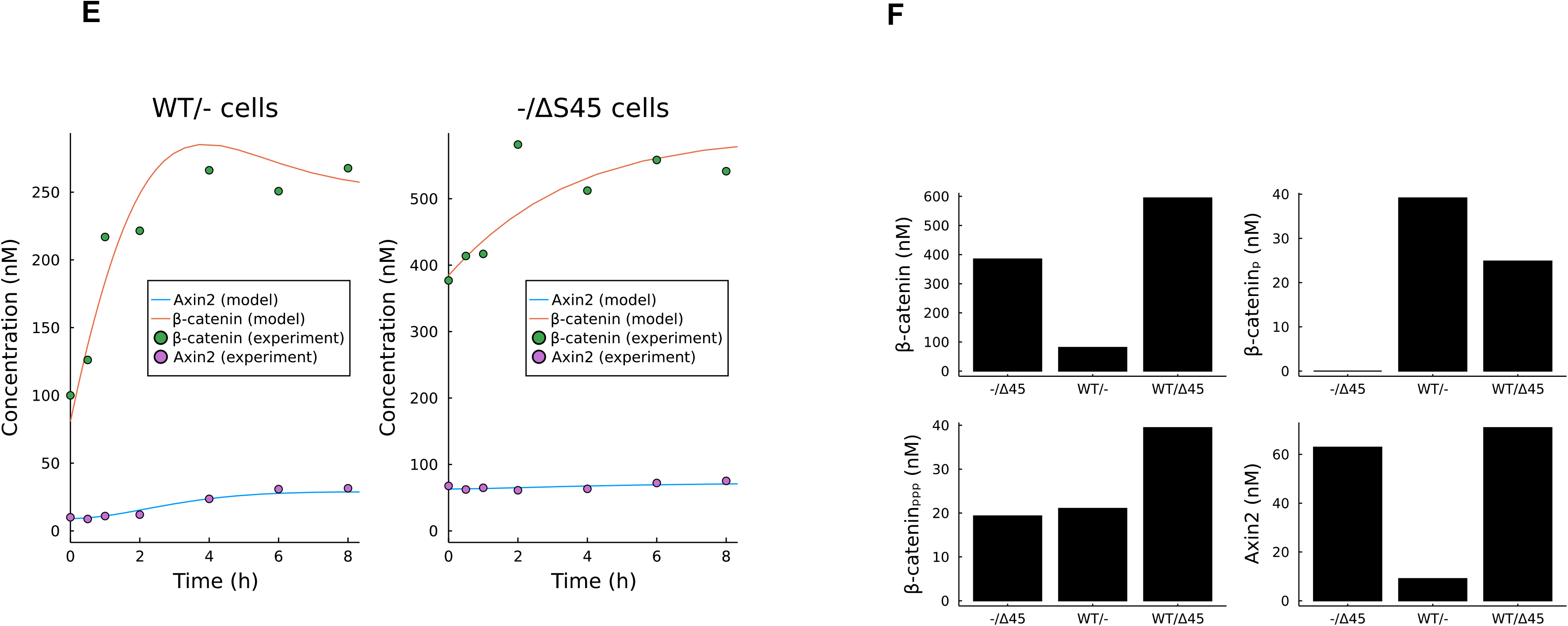
Negative feedback in ΔS45 β-catenin regulation. **A**, Scheme showing that in WT/ΔS45 cells, the protein concentration of WT β-catenin is predicted to drop in the presence of negative feedbacks. **B**, WT β-catenin protein concentration drops in WT/ΔS45 cells, as compared to WT/- cells, indicating the presence of negative feedbacks. WT β-catenin protein was detected using a specific phospho S45 β-catenin antibody. **C**, Phospho S45 β-catenin in untreated RKO cells lysates and in RKO cells lysates treated with λ-phosphatase. **D**, Expression of WT-myc-β-catenin and S45F-myc-β-catenin in 293T cells. **E**, Data fitting results for WT/- and -/Δ45 cells. See Supplemental Material for more information. **F**, Model prediction of the concentration of β-catenin, CK1α-phospho β-catenin, GSK3-phospho β-catenin, and Axin2 in WT/ΔS45, WT/- and -/ΔS45 cells.

### Quantitative modeling of the β-catenin dynamics in -/ΔS45 and WT/- cells

Our experiment has clearly shown that without S45 priming phosphorylation, β-catenin can still go through proteasome-dependent degradation promoted by GSK3 phosphorylation. However, it is hard to determine the contribution of different mechanisms because the experiment is reflecting a mixed output of different mechanisms. For example, both Axin1 and Axin2 can facilitate β-catenin degradation, and GSK3 phosphorylation could happen in both S45 dependent and independent manner. To better disentangle the different regulations, we have made a mathematical model based on our previous work (fig. S1).

Our finding of the S45-independent GSK3 phosphorylation, indicates there will be four different populations of β-catenin: *β* represents the non-phosphorylated population, *β_p_* the CK1-phosphorylated population, *β_ppp_* the GSK3-phosphorylated population, and *β_pppp_* the CK1 and GSK3-phosphorylated population. In the mutant cell, due to the deletion of S45 site on β-catenin, there are only two possible forms of β-catenin which is <u>are</u> denoted as *β_m_* and *β_mppp_*, representing the non-phosphorylated and GSK3-phosphorylated β-catenin, respectively. GSK3 works under two different scenarios, S45-phosphorylation dependent and independent ones. The first is denoted as rate constant *k_GSK_*, representing the S45<u>-</u>phosphorylation_ -dependent GSK3 activity. The second is denoted as *k_GSK’_*, representing the S45-phosphorylation independent GSK3 activity. The two rate constants are mathematically connected with a ratio factor.

By fitting the total β-catenin and Axin2 time course in both WT/- and -/Δ45 cells, we have determined all the parameters in the model (Supplement text, fig. 4E). By carefully looking into the parameter set, we could achieve a quantitative understanding of the weights of different regulation mechanisms. First, just by fitting the time courses, we conclude that the S45 priming-dependent GSK3 phosphorylation is about 53-fold more potent than the S45 independent ones. The fold change is similar to the 30-fold difference claimed by others (7). Second, we found that Wnt also regulates the S45--priming -independent GSK3 phosphorylation. This regulation is necessary to explain the change of β-catenin concentration change in -/Δ45 cells. However, this regulation is only 30% as strong as to the S45 priming-dependent phosphorylation. Third, Axin1 and Axin2 activity parameters show that, in the absence of Wnt ligand, the GSK3 phosphorylation from Axin1 is about 17-fold as strong as the Axin2 activity. Axin1 plays a dominant role, so Axin2 knock-out cells do not show clear β-catenin concentration change. But in the presence of Wnt ligand, the β-catenin phosphorylation activity from Axin2 is about 6-fold as strong as that from Axin1, showing the strong effect of Axin2 feedback in controlling the steady-state β-catenin concentration. Interestingly, our model can predict the correct order of the pS45 β-catenin phosphorylation (fig. 4F), where it is weakest in -/Δ45 cells, the next in WT/Δ45 cells, and the strongest in WT/- cells (fig. 4B).

## DISCUSSION

There are three important conclusions we wish to draw from this analysis of the classic and careful papers from the Vogelstein laboratory. First, there is no need to postulate that the basic biochemistry of the priming event on S45 by CK1α and the subsequent phosphorylations by the kinase GSK3 is <u>are</u> bypassed in cancer cells by some other process, or that there is a backup system that is somehow elicited in the tumor cell that allows the unprimed phosphorylations to take place. Rather, the unprimed phosphorylations are always there but very inefficient. The unprimed events in the tumor cells are simply due to mass action, caused by the rise in concentration of β-catenin, which in turn increases both Axin1 and Axin2. The more dramatic effect on Axin2 is of special interest. A recent study of β-catenin mutations in hepatocellular carcinoma (8, 9) concluded that the S45 mutation is weak unless accompanied by additional mutations in the third exon of β-catenin or APC.

While this may be true to some degree, this paper shows that they are not necessary to explain the GSK3 phosphorylations that are observed in the S45 mutations. We would argue that the existing negative feedback circuitry on the Axins fully explains these GSK3 phosphorylations in the S45 mutants. These experiments emphasize the importance of the Axin feedback loops in suppressing the effects of perturbations of β-catenin, whether or not they are caused by mutations in β-catenin and explain the conclusion that the S45 mutations seem weak. It may explain why most colorectal cancer mutations occur in APC, which is not directly inhibited by a rise in β-catenin. It is counter to expectations that ΔS45 mutations should be weak for cancer, whereas APC mutations should be strong. There is a general lesson here for interpreting the effects of mutations in vivo from even very high quality in vitro experiments, when the in vivo biochemical processes are imbedded in circuits. Although one can invent complex regulatory loops to explain the in vivo discrepancies, the answers may simply be due to mass action. We previously encountered the non-obviousness of mass action as an unavoidable and simple explanation for the dynamics of the Wnt system (2). Confusion arises when asking what additional process terminates the rise of β-catenin after Wnt stimulation, when the perfectly adequate answer is that as β-catenin accumulates the rate of degradation should increase just due to mass action. In fact, as we showed here, mass action offers a quantitatively adequate explanation. In the case of the S45 mutations and the persistent phosphorylation on the GSK3 sites seen in the S45 mutations we do not need to invoke novel feedback processes in the cells (such as additional mutations in other components) but it is not so simple that it can be attributed to the rise in β-catenin alone. Ultimately the level of phosphorylation in the presence of the S45 mutation is insufficient on its own, where degradation would balance the rate of synthesis at modest levels of β-catenin. However, there is another pathway that negatively controls β-catenin levels. Feedback loops of Axin, which are elicited by a rather small increase in β-catenin level, increases the level of Axin1 and Axin2 which effectively increases the degradation rate by mass action. These feedbacks on Axin1 and Axin2 probably offer many more points of regulation and are probably utilized to control β-catenin dependent transcription in different circumstances and in different cell types.

## METHODS

### Antibodies and reagents

The following antibodies were used: β-catenin (BD Transduction Laboratories, San Jose, CA), phospho-β-catenin (Ser45) (#2564; Cell Signaling, Danvers, MA), phospho-β-catenin (Ser33/37/Thr41) #9561; Cell Signaling), Axin1 (AF3287, R&D Sytems, Minneapolis, MN), Axin2 (#76G6; Cell Signaling), CK1α (AF4569, R&D Systems) and α-tubulin (NeoMarkers, Fremont, CA).

### Tissue culture and production of Wnt-3A conditioned media

Cell cultures were cultured in Dulbecco modified Eagle medium DMEM (RKO cells) or Mccoy’s 5A (HCT116 cells) containing 10% fetal calf serum and streptomycin-penicillin in 37°C humidified incubator with 5% CO_2_. The cell lines except for HCT116 (including WT/ΔS45, WT/- and -/ΔS45) were purchased from American Tissue Culture Collection (ATCC, Manassas, VA). WT/ΔS45, WT/- and -/ΔS45 cells were kindly provided by Bert Vogelstein. Wnt-3A conditioned media (Wnt-3A CM) and control media were prepared according to manufacturer’s protocol.

### Treatment with Wnt-3A and lithium chloride

Cultured cells were stimulated with Wnt-3A CM or 100 ng/ml recombinant mouse Wnt-3A (R&D Systems) for the indicated times. Cells were treated with 40 mM lithium chloride for one hour.

### Lentivirus-based transduction of small-hairpin RNA

The 19-base pair long shRNA sequences are aagaagatgtccacgcctg for human CK1α. A lentivirus based plasmid was used to deliver shRNAs to the target cells.

### Preparation of whole-cell protein lysates

The cells were washed twice with ice-cold PBS and lysed with a buffer containing 50 mM Tris/pH 7.6, 150 mM NaCl, 5 mM EDTA/pH 8.0, 0.5% NP-40, supplied with protease inhibitor cocktail (Roche), 1 mM phenylmethylsulfonyl fluoride, and the phosphatase inhibitors sodium fluoride (10 mM), p-nitrophenylphosphate (20 mM), β-glycerophosphate (20 mM), sodium orthovanadate (1 mM), okadaic acid (1 mM), and mycrocystin-LR (1 mM). The lysates were incubated on ice for 0.5 hour, and then cleared by full-speed centrifugation at 4°C for 0.5 hour.

The total protein concentration of the lysates was measured by the Bradford method (BioRad, Hercules, CA), using bovine serum albumin (BSA) (Sigma-Aldrich, St. Louis, MO) as a standard.

### Separation of membrane and cytoplasmic/nuclear β-catenin

In order to separate the membrane-associated and free β-catenin fractions, whole-cell lysates were incubated with Concanavalin A-Sepharose 4B (Sigma-Aldrich). Whole-cell lysates were rotated at 4°C for 1 hour with 100 ml of 50% Con-A slurry. The supernatant and the Con-A sepharose beads were separated by centrifugation (4°C, 5 minutes, 6,000X*g*). The Con-A bound protein fraction was eluted from the Con-A sepharose beads by incubating the beads with methyl-a-D-glucopyranoside (250 mM) and methyl-a-D-mannopyranoside (250 mM). The supernatant contains the free (cytoplasmic and nuclear) fractions of β-catenin, while the Con-A bound fraction contains the membrane-associated fraction of β-catenin.

### Qualitative and quantitative immunoblotting

Proteins were resolved by linear SDS-PAGE (7.5% acrylamide). The proteins were transferred into nitrocellulose membranes by electrophoretic transfer.

For qualitative immunoblotting, the nitrocellulose membranes were blocked with blocking buffer (TBS, 0.1% Tween-20, 5% nonfat dry milk) for 0.5 hour at room temperature. The membranes were incubated overnight at 4°C with a primary antibody, which was diluted 1:500 to 1:2000. After extensive washing with TBS/0.1% Tween-20, the membranes were incubated for 30 minutes at room temperature with a horseradish peroxidase (HRP)-conjugated secondary antibody (Jackson ImmunoResearch, West Grove, PA), which was diluted 1:10,000. The proteins were detected by Enhanced Chemiluminescence (ECL) using ECL Western Blotting reagents (Amersham, GE Healthcare, UK) or SuperSignal West Dura Extended Duration Substrate (Pierce, Rockford, IL, US). For quantitative immunoblotting, the nitrocellulose membranes were washed with PBS and blocked with Odyssey blocking buffer (Li-Cor, Lincoln, NE). The membranes were incubated overnight at 4°C with a primary antibody, which was diluted 1:500 to 1: 5,000. After extensive washing, the membranes were incubated for 1 hour at room temperature with a fluorescent secondary antibody, which was diluted 1: 10,000. The membranes were then scanned in the appropriate infrared channel using an Odyssey Infrared Scanning System (Li-Cor). The secondary antibodies used were Alexa Fluor 680 donkey anti-rabbit (Molecular Probes, Carlsbad, CA), and IRDye 800CW goat anti mouse (Li-Cor). The quantification of protein bands was performed using the Odyssey application software (Li-Cor). Protein quantification was calculated on base of measurements of at least three independent experiments in duplicates.

### RNA extraction, cDNA synthesis, and qRT-PCR

RNA was extracted from cells using RNeasy (Qiagen, Germany). First strand cDNA was synthesized from 0.5-1 mg RNA by Superscript III First-Strand Synthesis superMix for qRT-PCR (Invitrogen). qRT-PCR reactions were performed using Universal PCR master mix (Applied Biosystems, Branchburg, NJ) and TaqMan gene expression assays (Applied Biosystems), according to the manufacturer’s protocol. GAPDH and β-actin were used as endogenous controls, as the expression levels of these genes were found to remain unaffected by Wnt stimulation. Each sample was measured in triplicates. The qRT-PCR reactions were carried out in iCycle iQ (BioRad, Hercules, CA), as follows: 40 cycles of amplification with 15 second denaturation at 95°C and 1-minute annealing at 60°C. The gene expression analysis was done using iQ5 software (BioRad). Relative and normalized gene expressions were calculated (ΔΔCT). The gene expression was normalized to an endogenous control, such as GAPDH or β-actin, and was relative to a defined control condition.

