## Supplementary Materials for "The self-regulated response of the Wnt pathway to an oncogenic mutation in β-catenin"

**Supplement figure**

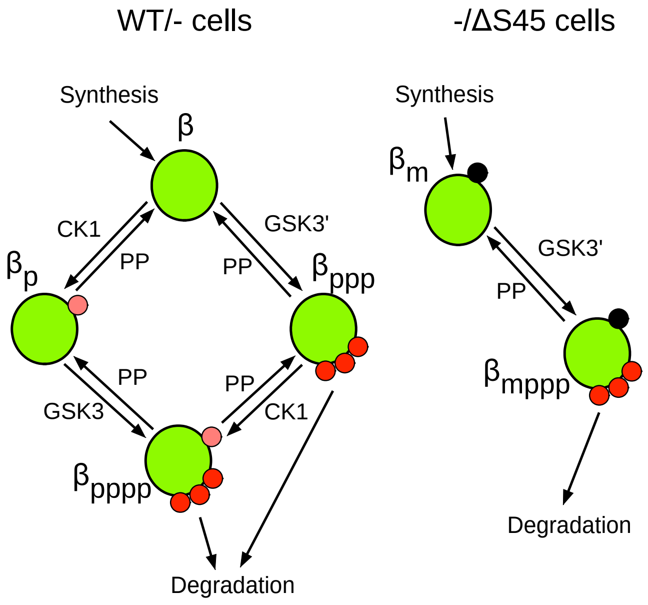


Figure S1: model diagram of detailed mass action regulations of different forms of β-catenin.

**Supplemental text**

The new model has the following assumptions:

1. There are four populations of β-catenin, $\beta$ is the non-phosphorylated population, $\beta_p$ is the CK1-phosphorylated population, $\beta_{ppp}$ is the GSK3-phosphorylated population, and $\beta_{pppp}$ is the CK1-and-GSK3-phosphorylated population,
2. CK1 activity is independent of 33/37/41 phosphorylation status of β-catenin
3. GSK3 activity has two fractions:
    a. GSK3 represents the CK1-priming dependent phosphorylation, which contains two parts: the Axin1-dependent-activity is regulated by Wnt pathway activity, whereas Axin2-dependent-activity is not. Mathematically, $k_{GSK} = k_1 + k_2 * Axin2$, where $k_1$ is the phosphorylation rate constant from Axin1 bound GSK3, and $k_2$ the phosphorylation rate constant from Axin2 bound GSK3.
    b. GSK3' represents the nonspecific GSK3 activity, which does not depend on the priming phosphorylation of CK1. This fraction, hoever, is dependent on the quantity of Axin1 and Axin2, practically, $k_{GSK3'} = GSK0Ratio * k_{GSK}$.
4. All the dephosphorylation rate constant are the same
5. Degradation speed is only dependent on the status of GSK phosphorylation sites, but not CK1 sites.
6. To mimic the Wnt pathway activation, we set a new value for $k_1$ and $k_{GSK0}$ at time 0.

Accordingly, we could write the rate equations as:

$$\frac{d}{dt}\begin{bmatrix}\beta\\ \beta_p\\ \beta_{ppp}\\ \beta_{pppp}\\ \beta_m\\ \beta_{mppp}\\ Axin2\end{bmatrix} = \begin{bmatrix} -k_{CK}-k_{GSK'} & k_p & k_p & 0 & 0 & 0 & 0\\ k_{CK} & -k_p-k_{GSK} & 0 & k_p & 0 & 0 & 0\\ k_{GSK'} & 0 & -k_p-k_{CK}-k_{rb} & k_p & 0 & 0 & 0\\ 0 & k_{GSK} & k_{CK} & -2k_p-k_{rb} & 0 & 0 & 0\\ 0 & 0 & 0 & 0 & -k_{GSK'} & k_p & 0\\ 0 & 0 & 0 & 0 & k_{GSK'} & -k_p-k_{rb} & 0\\ 0 & 0 & 0 & 0 & 0 & 0 & -k_{ra2}\end{bmatrix}\begin{bmatrix}\beta\\ \beta_p\\ \beta_{ppp}\\ \beta_{pppp}\\ \beta_m\\ \beta_{mppp}\\ Axin2\end{bmatrix} + \begin{bmatrix}k_S\\ 0\\ 0\\ 0\\ k_{sm}\\ 0\\ k_{a02}+k_{a2}Hill(\beta_{total})\end{bmatrix}$$

We use Julia language to code and simulate the Axin2 model. Most of the parameters in the model are not experimentally measured. To determine the parameter values, we use the function "DiffEqFlux.sciml_train" to fit the model to experimental time-course data (Fig. 3E and Fig. 4A). One key element to have a good fitting is the loss function. The loss function we used here is:

$$
\begin{aligned}
Loss = & \sum_{n=all\ time\ points} \left(\frac{\beta_{sim}}{\beta_{exp}} - 1\right)^2 + \sum_{n=all\ time\ points} \left(\frac{\beta_{exp}}{\beta_{sim}} - 1\right)^2 \\
& + \sum_{n=all\ time\ points} \left(\frac{Axin2_{sim}}{Axin2_{exp}} - 1\right)^2 + \sum_{n=all\ time\ points} \left(\frac{Axin2_{sim}}{Axin2_{exp}} - 1\right)^2 \\
& + \sum_{n=all\ time\ points} \left(\frac{\beta_{p,sim}}{\beta_{p,sim}(t=0)} - \beta_{p,exp}\right)^2 \\
& + \sum_{n=all\ time\ points} \left(\frac{\beta_{ppp,sim}}{\beta_{ppp,sim}(t=0)} - \beta_{ppp,exp}\right)^2
\end{aligned}
$$

The first four terms are used to minimize the difference between the measured and simulated β-catenin and Axin2 concentrations. We set the β-catenin concentration at time 0 in the WT/- cell to be 100 nM. And the Axin2 concentration to be 10nM. These are estimations in the reasonable range. The exact concentration in the WT/- cell line is unknown. The last two terms minimize the difference between the measured and simulated CK1-phosphorylated and GSK-phosphorylated β-catenin. Because we do not have the measurement for phosphorylated β-catenin concentration in any cell line, we fitted the normalized concentration.

| Parameter | Value | Explanation |
|---|---|---|
| $k_s$ | 4.97 nM/min | β-catenin translation rate |
| $k_{rb}$ | 0.116 $min^{-1}$ | Degradation rate constant for GSK3 phosphorylated β-catenin |
| $k_{ck}$ | 2.4 $min^{-1}$ | The rate constant of CK1 phosphorylation |
| $k_p$ | 2.53 $min^{-1}$ | The rate constant of dephosphorylation |
| $k_{a02}$ | 0.042 nM/min | Nonregulated endogenous Axin2 translation rate constant |
| $k_{a2}$ | 0.445 nM/min | The rate constant of regulated Axin2 translation |
| $k_{ra2}$ | 0.00541 $min^{-1}$ | The rate constant of Axin2 degradation |
| $K$ | 200.5 nM | Hill coefficient for the Hill function of Axin2 activation |
| $k_1$ | 2.03 nM/min | Axin1 dependent and CK1-priming dependent GSK3 phosphorylation rate constant |
| $k_2$ | 0.0123 $min^{-1}$ | Axin2 dependent and CK1-priming dependent GSK3 phosphorylation rate constant |
| *GSK0Ratio* | 0.0187 | Relative GSK3 activity of non-CK1 primed / CK1 primed |
| *Ck1fold* | 1.1 | Fold change of $k_{ck}$ upon Wnt pathway activation |
| *Gskfold* | 16.8 | Fold change of $k_{gsk}$ upon Wnt pathway activation |
| *Gsk0fold* | 0.288 | Fold change of *GSK0Ratio* upon Wnt pathway activation |

From the fitted parameters, we have the following conclusions:

1. Gsk0fold is less than 1, meaning the nonspecific GSK3 phosphorylation of β-catenin is also Wnt pathway regulated. The regulation is necessary to explain the slight increase of β-catenin in -/Δ45 cells.
2. *GSK0Ratio* represents the activity difference between the CK1-independent GSK phosphorylation and the CK1-dependent GSK3 phosphorylation, which is about 53-fold difference.
3. The activity that Axin2 degrade $\beta$ ($Axin2*k_2$) is about 0.123 nM/min. The activity from Axin1 is about 17-fold as fast as the Axin2 regulation. After the Wnt3a activation, the activity from Axin2 becomes 0.775 nM/min due to the increased Axin2 concentration, while the activity from Axin1 becomes 0.12 nM/min due to the Wnt pathway regulation, more than 6-fold less significant than Axin2. This means that without Wnt pathway activation, the contribution of β-catenin degradation from Axin2 is negligible, while Axin2 becomes substantially more critical after the pathway activation.
